# Decoding plant x environment interactions through spatial gradients in leaf physiology and metabolism

**DOI:** 10.64898/2026.10.14.759099

**Authors:** Rubén Pérez Valverde, Thomas Nägele

**Author notes:** Correspondence, T.

## Abstract

Succulent leaves are well-known to combine photosynthesis, carbon storage and hydraulic buffering. However, their metabolic responses to heat and drought may differ along the same leaf axis. We investigated spatial and temporal changes in leaf tissue water content, photosynthetic performance and central carbon metabolism in the CAM plant *Sansevieria cylindrica* across three experimental phases: (i) heat, (ii) combined heat and drought, and (iii) rewatering under continued heat. Apical, middle and basal leaf sections were sampled at midday, and chlorophyll fluorescence was monitored near the leaf tip. Fresh-weight-to-dry-weight ratios declined significantly in basal tissues under drought, whereas apical tissues showed a significant decrease only during rewatering indicating a considerable delay in tissue water supply along the leaf axis. Apical photosystem II quantum yield decreased during combined heat and drought, and sucrose accumulated transiently during heat exposure, particularly in middle and basal sections. Glucose and fructose peaked under drought conditions and declined after rewatering. Citrate decreased during the experimental time course, while apical malate increased significantly during rewatering phase. Chlorophyll varied primarily with leaf position, whereas starch and fumarate showed no significant effects. Neural network classification identified citrate and sucrose as major contributors to the discrimination of treatment phases, while chlorophyll was most informative for distinguishing leaf positions. Together, these findings demonstrate that within-leaf physiological and metabolic gradients capture complementary dimensions of plant–environment interactions, distinguishing spatially structured leaf characteristics from dynamic stress responses. Integrating spatially resolved metabolic and physiological profiles thus provides a promising approach for identifying biochemical signatures of environmental stress and recovery in CAM plants.

## Introduction

Drought represents one of the major environmental constraints on plant growth, productivity, and geographical distribution (Craine et al., 2013; Brodrick et al., 2019). The capacity of plants to withstand prolonged water limitation depends on coordinated adjustments at morphological, physiological, and metabolic levels (Gupta et al., 2020). Among these responses, the regulation of carbon metabolism is particularly important because drought alters photosynthetic carbon assimilation while simultaneously changing the demand for carbon skeletons, energy, osmotic adjustment, and cellular maintenance. Soluble carbohydrates and organic acids play central roles in these processes. In addition to their roles as substrates and intermediates of primary metabolism, these compounds contribute to osmotic regulation, redox balance, carbon storage, and the coordination of photosynthetic and respiratory metabolism (Geigenberger et al., 2005). Characterizing their spatial distribution can therefore provide insight into how drought-adapted plants regulate carbon metabolism under conditions in which water conservation strongly constrains gas exchange.

Succulent species provide particularly informative systems in which to investigate these mechanisms. The monocotyledonous genus *Sansevieria* (Asparagaceae) comprises approximately 70 species distributed across Africa and Asia, with East Africa representing the principal centre of species diversity (Brown, 1915; Webb and Newton, 2017). A characteristic feature shared by all members of the genus is the distinct differentiation of the mesophyll into chlorenchyma and a specialized central, colourless tissue devoted to water storage (Koller and Rost, 1988). In addition to dispersed vascular and fibre bundles, this water-storage tissue consists of two distinct cell types. Large, thin-walled, non-living water-storage cells occur in clusters and serve as the primary sites of water accumulation (Alfani et al., 1989). These clusters are surrounded and interconnected by a network of smaller living cells, referred to as network cells (Koller and Rost, 1988). Although this basic anatomical organization appears to be conserved throughout the genus, the relative proportions of the two cell types differ among species. Considerable interspecific variation also occurs in leaf morphology, xeromorphic adaptations, and the extent of succulence. Leaves may be cylindrical or relatively flat and concave, while succulence can range from hydrenchyma-based water storage to all-cell succulence, with substantial differences in the degree of succulence among species (Alfani et al., 1989).

*Sansevieria cylindrica* is a drought-adapted succulent characterized by cylindrical leaves, substantial internal water-storage capacity, and crassulacean acid metabolism (CAM; Martin et al., 2019). CAM represents an important adaptation to environments in which water availability limits daytime gas exchange. In CAM plants, atmospheric CO₂ is predominantly fixed at night by phosphoenolpyruvate carboxylase, producing organic acids, e.g. malate, that are accumulated in the vacuole. During the subsequent light period, these acids are decarboxylated, supplying CO₂ internally to the Calvin-Benson-Bassham cycle while stomatal opening and water loss by transpiration remain low. This temporal separation of initial CO₂ fixation from photosynthetic carbon reduction confers high water-use efficiency but requires tight coordination between carbohydrate turnover, organic acid metabolism, vacuolar storage, and the circadian regulation of carbon fluxes (Borland and Taybi, 2004; Borland et al., 2016).

Although CAM is commonly characterized through temporal changes in gas exchange, acidity, and metabolite concentrations, metabolic regulation may also vary substantially across a leaf blade. Differences in tissue age, developmental state, source–sink relations, hydraulic status, and local photosynthetic capacity can generate metabolic gradients along the leaf axis (Smith et al., 1997). Such heterogeneity may be especially pronounced in long-lived succulent leaves, in which tissues at the base and tip differ in developmental history and potentially in their exposure to water limitation. Measurements based exclusively on whole-leaf material can obscure these differences by averaging metabolically distinct regions and may consequently conceal regulatory patterns that are important for understanding drought adaptation.

Spatially resolved analysis within a single leaf offers several advantages for investigating these processes. Most importantly, comparing positions along the same leaf reduces variation associated with differences among individual plants, leaves, developmental stages, and environmental histories. The resulting metabolic gradient can reveal whether carbohydrate reserves, soluble sugars, and organic acids are distributed uniformly or whether distinct regions preferentially perform particular metabolic functions. Because carbohydrates provide substrates for nocturnal phosphoenolpyruvate regeneration in CAM, while organic acids represent major products and transient carbon stores of nocturnal fixation, coordinated gradients in these metabolite classes may indicate spatial differences in CAM activity and carbon allocation. Conversely, divergent distributions could point to differences in storage capacity, respiratory demand, source–sink function, or local responses to water status.

Understanding such within-leaf organization is particularly relevant to drought tolerance. Succulent leaves simultaneously function as photosynthetic organs, water reservoirs, and sites of carbon storage, creating the possibility that progressive water limitation is accompanied by spatial reorganization of metabolism rather than a uniform response throughout the organ. Local differences in water availability could alter carbohydrate mobilization and organic acid turnover, allowing particular regions to maintain metabolic activity or osmotic balance as drought progresses. Resolving these patterns may therefore provide information that can hardly be obtained from whole-leaf measurements and may help distinguish constitutive spatial organization from metabolic adjustments associated with drought adaptation.

In the present study, we investigated the distribution of carbohydrates and organic acids along the leaf blade of *S. cylindrica* to determine whether the central carbon metabolism exhibits a systematic gradient. By comparing metabolite dynamics together with photosynthetic performance during heat and drought exposure among top, middle and bottom leaf positions, we aimed to characterize spatial coordination between carbohydrate availability and organic acid metabolism and to assess how such organization may contribute to the regulation of metabolism. This within-organ perspective provides a framework for examining metabolic adjustments during heat and drought exposure not only as a whole-plant response, but also as a spatially structured metabolic process occurring within a single photosynthetic organ.

## Materials and Methods

### Plant material and growth conditions

Plants of *Sansevieria cylindrica (‘Mikado’*) with leaf length of 15–20 cm were purchased (www.pflanzmich.de) and acclimated for two months at 22–25 °C during the day and 18–20 °C at night under a 12 h light/12 h dark cycle. Photosynthetic photon flux density was 200–250 µmol photons m⁻² s⁻¹ at the leaf top, 100–150 µmol photons m⁻² s⁻¹ at the middle and 50–80 µmol photons m⁻² s⁻¹ at the base. Apical, middle and basal regions were defined as equal thirds of the aboveground leaf length (Supplementary Figure SF1). Plants were watered weekly before the experiment. Water was then withheld until sampling on day 8, after which plants were rewatered until day 12. Temperatures of 30–33 °C during the light period and 20–22 °C at night were maintained throughout the experimental period, including rewatering. Relative humidity ranged from approximately 15% during the day to 60% at night. Air temperature, relative humidity and soil moisture were recorded using an Arduino UNO R4® WiFi platform equipped with a DHT22 temperature/humidity sensor and a Funduino® soil moisture sensor (Supplementary Figures SF2 and SF3). Day 0 served as the initial reference sampling point. A graphical summary of the experimental setup is provided in the supplements (Supplementary Figure SF4).

### Chlorophyll fluorescence measurements by pulse amplitude modulation fluorometry

The effective quantum yield of photosystem II, Y(II), was recorded throughout the experiment using a JUNIOR-PAM fluorometer (Heinz Walz GmbH, Effeltrich, Germany) operated with the WinControl® software. The fibre-optic probe was positioned 2 cm below the tip of a randomly selected leaf, at a fixed distance and angle, and remained in place throughout the experiment. The complementary yields of regulated, Y(NPQ), and non-regulated, Y(NO), energy dissipation were determined according to Kramer et al. (2004), such that Y(II) + Y(NPQ) + Y(NO) = 1. Daytime and nighttime measurements were evaluated separately.

### Leaf sampling

Leaf material was harvested at midday on days 0, 2, 6, 8 and 12, covering the initial reference (0d), water-deprivation (2-8d) and rewatering (8-12d) phases under continued heat. Leaves were cut with a blade and divided into apical, middle and basal thirds, each approximately 5–6 cm long. Leaf discs were weighed to determine fresh weight, and material for metabolite analysis was extracted immediately as described below. For fresh weight to dry weight ratios, leaf discs collected on days 0, 8 and 12 were weighed fresh and dried at 70 °C for three days before determination of dry weight (n = 5 per group, as reported in Figure 1).

**Figure 1.**
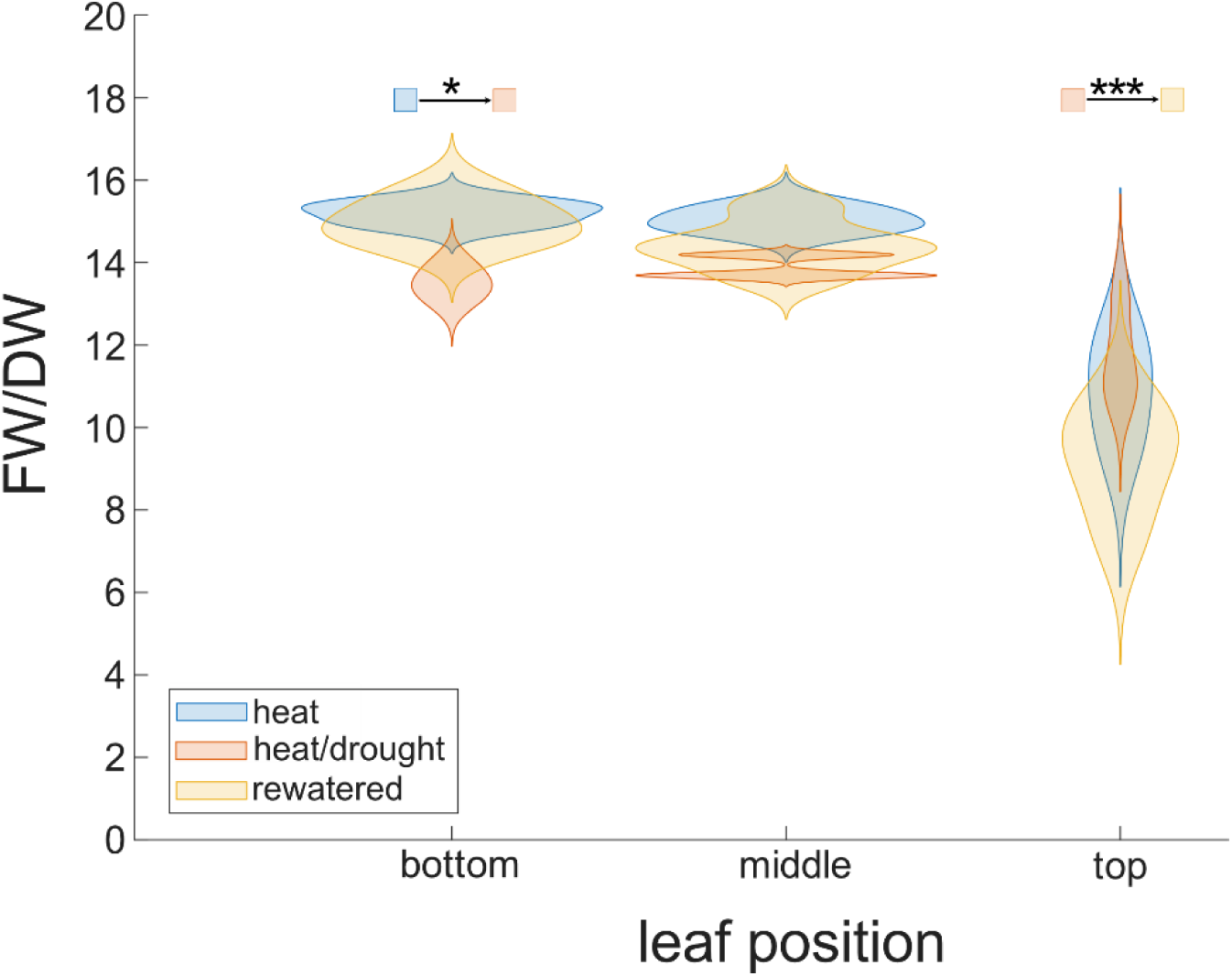
Fresh weight to dry weight ratios at different leaf positions during drought and recovery phases. To determine the ratios, leaf discs were cut at the different leaf positions, weighed, and then dried at 70 °C for 3 days before the dry weight was determined. Initial heat (blue violins): 0d; heat/drought (red violins): after 8d of water deprivation; rewatered (yellow violins): after 4 days of rewatering (in total: day 12 of the experimental time course). Asterisks indicate significance for comparisons indicated by coloured squares (ANOVA + Tukey’s post hoc test; *: p < 0.05; *** p < 0.001; n = 5 from 3 leaves).

### Chlorophyll quantification

Chlorophyll was extracted and quantified by the high-temperature ethanol method following established protocols (Palta, 1990). Weighed leaf fragments were homogenised directly in 400 µl 80% (v/v) ethanol together with a small amount of sea sand to aid mechanical disruption, incubated at 80°C for 30 min in sealed tubes and centrifuged for 10 min to sediment cellular debris and sand. The supernatant, containing pigments and soluble carbohydrates, was transferred to a fresh tube, and the ethanol extraction was repeated on the pellet. Afterwards, the two supernatants were pooled.

A 200 µl aliquot of the pooled ethanol extract was transferred to a 96-well plate and absorbance was read at 654 nm, using 80% (v/v) ethanol as the blank. Total chlorophyll content (TCC, µg ml−1) was calculated as TCC = 1000 × A_654_/39.8 and normalised to the fresh weight of the corresponding leaf segment. The remaining ethanol extract was retained for soluble carbohydrate quantification. Starch was determined in the pellet of the ethanol extraction.

### Soluble carbohydrate measurements

The pooled ethanol extracts described above were evaporated to dryness in a desiccator, then resuspended in 500 µl H_2_O and shaken at 750 rpm for 45 min at ambient temperature. Samples were centrifuged at 13 000 rpm for 15 min and the supernatants were transferred to fresh tubes for sucrose, glucose and fructose quantification. For samples which exceeded the linear range of the respective calibration curve, extracts were diluted in H_2_O by factors of 1:20 and 1:40.

Sucrose was quantified by the anthrone assay as described before (Brodsky et al., 2026). A 100 µl aliquot of extract, or of sucrose calibration standard, was mixed with 100 µl 30% (w/v) KOH and incubated for 10 min at 95°C to destroy reducing sugars, leaving the non-reducing sucrose intact, and then cooled to ambient temperature.

Anthrone reagent (1 ml; 0.14% (w/v) anthrone in 14.6 M H_2_SO_4_) was added, the mixture was incubated at 40°C for 30 min, and absorbance was read at 620 nm. Sucrose contents were calculated against a calibration curve of authentic sucrose in H_2_O over the range 0–5 mM, processed identically.

Glucose was quantified in a coupled enzymatic assay in 96-well microplate format, following NADPH formation at 340 nm. A 20 µl aliquot of extract was incubated with 100 µl reaction buffer containing 100 mM imidazole-HCl (pH 6.9), 2 mM MgCl_2_, 2 mM ATP, 1 mM NADP^+^ and glucose 6-phosphate dehydrogenase (G6PDH, 1 U µl^−1^; 1 µl ml^−1^ buffer). Absorbance at 340 nm was recorded until stable to establish the background. Hexokinase (1 µl, 0.5 U µl^−1^) was then added, phosphorylating glucose to glucose 6-phosphate, which is oxidised by G6PDH with concomitant reduction of NADP^+^, and absorbance was followed to a stable endpoint. Glucose contents were calculated from the background-corrected endpoint against a calibration curve of glucose in H_2_O over the range 0–1 mM.

Fructose was quantified in the same well immediately afterwards. Phosphoglucose isomerase (1 µl, 2.5 U µl^−1^) was added, converting the fructose 6-phosphate formed from fructose by hexokinase into glucose 6-phosphate, which enters the same G6PDH reaction, and absorbance at 340 nm was again followed to a stable endpoint. The fructose signal was obtained as the difference between the second and the first endpoint, and fructose contents were calculated against a calibration curve of fructose in H_2_O over the range 0–1 mM.

### Starch quantification

Starch was quantified following Brodsky et al. (2026). After ethanol extraction, the insoluble pellet was treated with 0.5 N NaOH at 95 °C for 60 min, acidified with 1 M acetic acid and digested with amyloglucosidase. Released glucose was measured at 340 nm using a coupled hexokinase/glucose 6-phosphate dehydrogenase assay, following NADPH formation. Starch concentrations were normalised to tissue fresh weight and expressed as C6 equivalents.

### Quantification of organic acids

Carboxylic acids were extracted by adding double-distilled water preheated to 60 °C to dried plant material, followed immediately by incubation at 95 °C for 15 min. After brief centrifugation, the supernatant was analysed photometrically according to the manufacturer’s instructions. Carboxylic acid contents were determined in three independent biological replicates. Malate and citrate were quantified using the Malate Assay Kit (MAK511) and the Citrate Assay Kit (MAK333) (Sigma-Aldrich, St Louis, MO, USA) following the manufacturer’s instructions, with absorbance read at 565 nm. Fumarate was quantified using the Fumarate Assay Kit (MAK060) (Sigma-Aldrich) following the manufacturer’s instructions, with absorbance read at 450 nm.

### Statistical analysis, data visualization and neural network classification

Analyses and figures were produced in R using RStudio v.1.1. Chlorophyll and metabolite concentrations were analysed by two-way ANOVA with sampling day, leaf position and their interaction as factors, followed by Tukey–Kramer comparisons. Fresh weight to dry weight ratios were compared by ANOVA followed by Tukey’s post hoc test. The significance threshold was α = 0.05. Boxplots show medians, interquartile ranges (IQRs) and whiskers extending to the most extreme observations within 1.5 × IQR of the box; points beyond the whiskers denote outliers. Results of ANOVA and post-hoc tests is provided in the supplements (Supplementary Tables ST3 and ST4).

Classification followed the two-layer feedforward approach described before (Brodsky et al. 2026), implemented in MATLAB R2025b with the Deep Learning Toolbox. Sigmoid activation in the hidden layer captured nonlinear relationships, and a softmax output layer estimated the class probabilities. Data was partitioned into training (70%), validation (15%) and test (15%) subsets. Separate models classified treatment phase (initial reference, heat/drought and rewatering), sampling day (0, 2, 6, 8 and 12) and leaf position (top, middle and bottom). Class-specific receiver operating characteristic curves and AUCs were evaluated on the test subset using a one-versus-rest approach. ANOVA-based feature scores were calculated as −log(p), with larger values indicating stronger univariate associations with class labels.

## Results

To determine whether leaf water content varied across the leaf blade, leaf discs were collected from different positions along the leaf, and their fresh weight (FW) was compared with their dry weight (DW; Figure 1). Drought significantly affected the FW/DW ratio only in tissue from the basal region of the leaf (ANOVA, p < 0.05). After eight days without watering, the FW/DW ratio of basal leaf tissue decreased from 14–16 to 12–14. A similar but statistically non-significant decrease was observed in tissue from the middle of the leaf, whereas no effect was detected in tissue from the top of the leaves. While rewatering resulted in a reversed shift of FW/DW towards values of initial heat samples in bottom and middle positions, leaf tissue at the top showed a significant decrease of FW/DW ratios (ANOVA, p < 0.001). These observations demonstrated that the timing of changes in FW/DW differed along the leaf axis, with a delayed decrease in apical tissues after rewatering under continued heat.

Photosynthetic performance was monitored during day and night of each sampling time point (Figure 2). Daily medians of the effective quantum yield of photosystem II, Y(II), showed a significant decrease between day 6 and 8, while medians of night measurements continuously decreased from the beginning of the experiment (Figure 2 A). Watering after 8 days resulted in a slight increase of Y(II) during day and night until day 12. An opposite trend was observed for the yield of non-regulated energy dissipation, Y(NO), which increased during the time course both during the day and night (Figure 2 B). Non-significant fluctuations in the yield of (regulated) non-photochemical quenching, Y(NPQ), were observed in both daytime and nighttime measurements throughout the experimental time course (Figure 2C).

**Figure 2.**
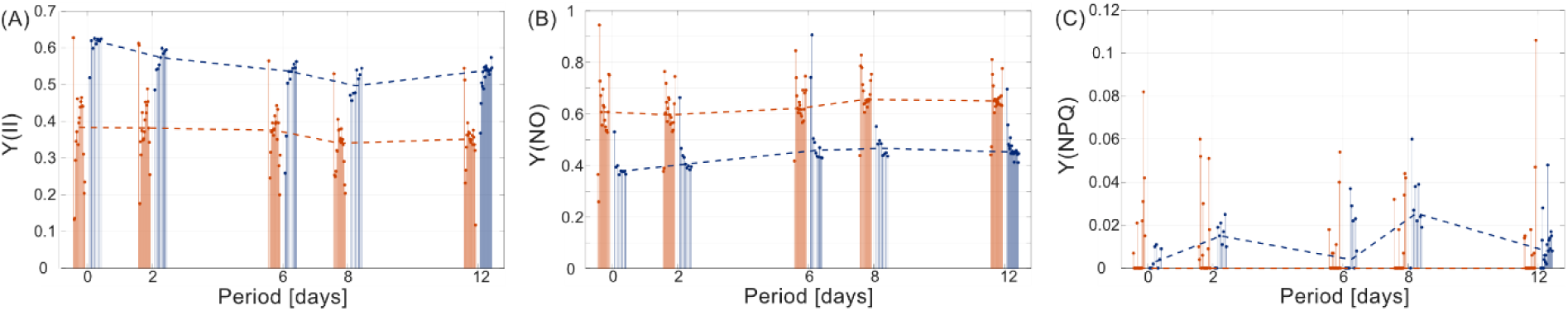
Time course of photosynthetic efficiency and non-photochemical quenching. **(A)** Effective quantum yield of photosystem II. **(B)** Yield of non-regulated non-photochemical quenching. **(C)** Yield of regulated non-photochemical quenching. Stems represent measurements during the experimental time course (orange: day measurements; blue: night measurements). Dashed lines represent linear interpolations of medians of the respective full day or night data set. All data is provided in the supplements (Supplementary Table ST1).

Pearson correlation revealed a significant negative correlation between Y(II) and Y(NO) both for daytime and nighttime measurements (Figure 3 A, C). Particularly for nighttime measurements, a separation of data derived during initial, stress and recovery phases could be observed with lowest Y(NO) values during the initial experimental phase (Figure 3 C, blue dots). Although being significant during the daytime, correlations between Y(II) and Y(NPQ) were much weaker both during day and nighttime (Figure 3 B, D). Overall, these results indicate a strong inverse relationship between Y(II) and Y(NO), whereas Y(NPQ) showed only a weak association with changes in Y(II), irrespective of the measurement period.

**Figure 3.**
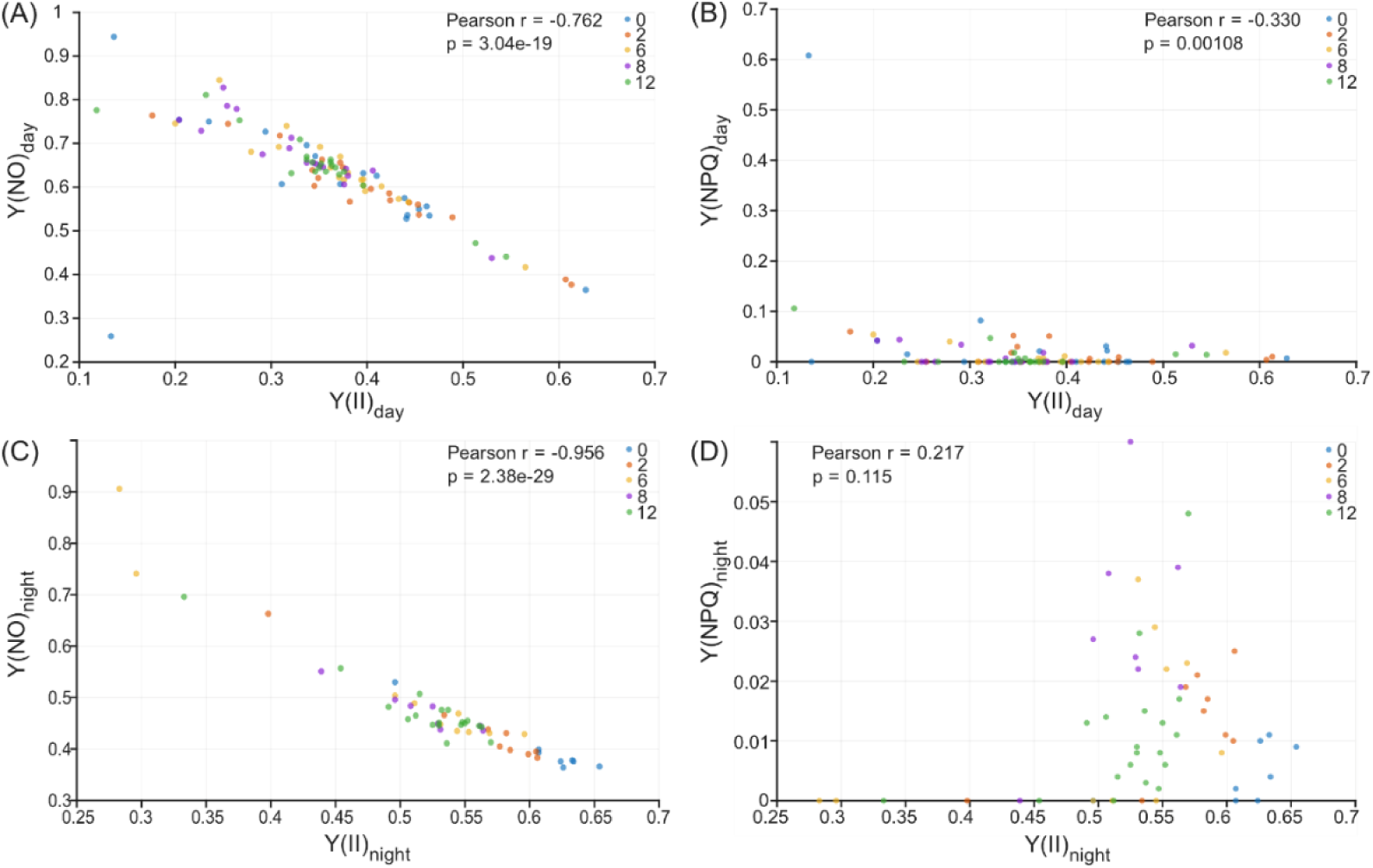
Relation between effective quantum yield of photosystem II and non-photochemical quenching. **(A)** Scatter plot of Y(NO) and Y(II) determined in day measurements. **(B)** Scatter plot of Y(NPQ) and Y(II) determined in day measurements. **(C)** Scatter plot of Y(NO) and Y(II) determined in night measurements. **(D)** Scatter plot of Y(NPQ) and Y(II) determined in night measurements. Colours of data points indicate time points of the experimental time course. Blue: 0 days; orange: 2 days; yellow: 6 days; purple: 8 days; green: 12 days. For each data pairing the Pearson correlation coefficient r is provided together with the p-value of the correlation.

To investigate temporal and spatial changes in (carbon) metabolism, the concentrations of chlorophyll, soluble sugars, and organic acids were determined in leaves collected from top, middle, and bottom positions over the experimental time course of 12 days (Figure 4). Two-way analysis of variance (ANOVA) was used to assess the effects of sampling time, leaf position, and their interaction. Significant effects were further examined using Tukey–Kramer post-hoc comparisons.

**Figure 4.**
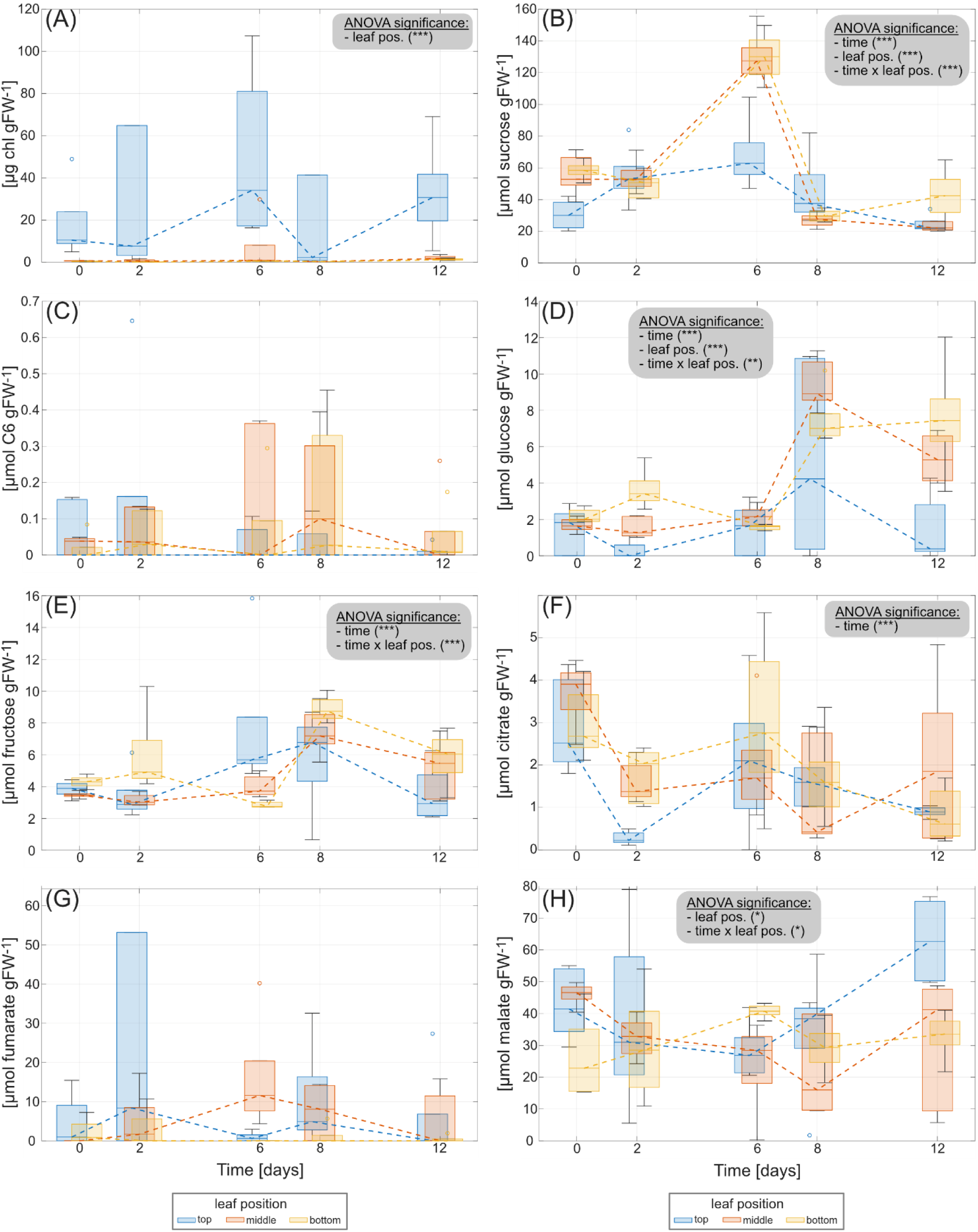
Temporal changes in metabolism across different leaf positions. Changes in the concentrations of (**A**) chlorophyll, (**B**) sucrose, (**C**) starch, (**D**) glucose, (**E**) fructose, (**F**) citrate, (**G**) fumarate, and (**H**) malate over the experimental time course sampled from three positions: top (blue), middle (orange), and bottom (yellow). Concentrations are expressed as µg g⁻¹ fresh weight (FW) for chlorophyll and µmol g⁻¹ FW for all other metabolites. Boxplots illustrate the distribution of measured concentrations at each sampling time (days 0, 2, 6, 8, and 12), with horizontal lines indicating medians, boxes representing interquartile ranges, whiskers showing data spread, and circles indicating outliers. Dashed lines connect median values across sampling times to illustrate temporal trends. The effects of sampling time, leaf position, and their interaction were evaluated using two-way analysis of variance (ANOVA). Significant effects are indicated within individual panels: P < 0.05 (*), P < 0.01 (**), and P < 0.001 (***). Only statistically significant effects are shown. All data is provided in the supplements (Supplementary Table ST2).

Chlorophyll concentration differed significantly among leaf positions (ANOVA, p < 0.001; Figure 4 A), whereas no significant effect of sampling time or interaction between sampling time and leaf position was detected. Throughout the experiment, chlorophyll concentrations were consistently higher in the apical leaf sections than in the middle and basal leaf sections. Post-hoc comparisons confirmed significantly higher chlorophyll concentrations in the apical leaf sections compared with both middle (p < 0.001) and basal leaf sections (p < 0.001). In contrast, chlorophyll concentrations did not differ significantly between the middle and basal leaf sections. Although chlorophyll concentrations in the apical leaf sections fluctuated over the experimental period, no significant temporal differences were identified. These findings indicate that chlorophyll concentration was primarily associated with leaf position rather than sampling time.

Soluble sugar concentrations showed pronounced temporal changes, with distinct accumulation patterns for sucrose, glucose, and fructose. Sucrose concentrations were significantly affected by sampling time (ANOVA, p < 0.01), leaf position (p < 0.001), and their interaction (p < 0.001; Figure 4 B). A pronounced increase in sucrose concentration was observed on day 6, particularly in the middle and basal leaf sections. Post-hoc analysis confirmed that sucrose concentrations on day 6 were significantly higher than those on all other sampling days (p < 0.001). Furthermore, sucrose concentrations were significantly higher in the middle (p = 0.0014) and basal leaf sections (p < 0.001) compared with the apical leaf sections when averaged across sampling times, whereas no significant difference was observed between middle and basal leaf sections. The significant interaction between sampling time and leaf position was reflected in the particularly pronounced accumulation of sucrose in the middle and basal leaf sections on day 6. At this sampling time, sucrose concentrations in both positions were significantly higher than those in the apical leaf sections (both p < 0.001). Subsequently, sucrose concentrations decreased markedly by day 8 and remained comparatively low until day 12. No significant difference was detected between days 8 and 12.

Starch amounts remained at low levels throughout the experiment (Figure 4 C). Although some fluctuations were observed, particularly in the middle and basal leaf sections, no statistically significant effects of sampling time, leaf position, or their interaction were identified.

Glucose concentrations were significantly affected by sampling time (ANOVA, p < 0.001), leaf position (p < 0.001), and their interaction (p < 0.01; Figure 4 D). Glucose concentrations remained relatively low during the initial sampling period but increased markedly on day 8, particularly in the middle and basal leaf sections. Post-hoc comparisons confirmed that glucose concentrations on day 8 were significantly higher than those on days 0, 2, and 6 (all p < 0.001). Although concentrations decreased by day 12, they remained significantly higher than those measured on days 0, 2, and 6 (all p < 0.001). The decrease between days 8 and 12 was also statistically significant (p = 0.0029). Across sampling times, glucose concentrations were significantly higher in the middle (p < 0.001) and basal leaf sections (p < 0.001) than in the apical leaf sections, whereas no significant difference was detected between middle and basal leaf sections. The significant interaction was particularly evident on day 8, when glucose concentrations in the middle leaf sections were higher than those in the apical leaf sections at the same sampling time (p < 0.001).

Fructose concentrations were significantly affected by sampling time (ANOVA, p < 0.001) and the interaction between sampling time and leaf position (p < 0.001; Figure 4 E). Fructose concentrations increased during the experiment, reaching their highest values on day 8. Post-hoc comparisons demonstrated that fructose concentrations on day 8 were significantly higher than those on days 0 (p < 0.001), 2 (p < 0.001), and 6 (p = 0.0033). Fructose concentrations subsequently declined significantly between days 8 and 12 (p = 0.0029). The significant interaction between sampling time and leaf position was reflected in differences in the magnitude of fructose accumulation. In particular, fructose concentrations at the basal leaf sections on day 8 were significantly higher than those measured in the same leaf position on days 0 (p = 0.0196) and 6 (p < 0.001). Similarly, fructose concentrations in the middle section of the leaves were significantly higher on day 8 than on day 2 (p = 0.0349).

The organic acids citrate, fumarate, and malate exhibited different patterns of accumulation throughout the experiment (Figure 4 F–H). Citrate concentrations were found to be significantly influenced by sampling time (ANOVA, p < 0.001; Figure 4 F). Concentrations were relatively high at the beginning of the experiment and subsequently decreased. Post-hoc comparisons confirmed significantly higher citrate concentrations on day 0 compared with days 2 (p < 0.001), 8 (p = 0.0012), and 12 (p < 0.001). In contrast, no significant difference was detected between days 0 and 6, indicating a transient recovery of citrate concentrations around day 6. No significant main effect of leaf position was observed.

Fumarate concentrations exhibited considerable variability among sampling times and leaf positions (Figure 4 G). Although elevated concentrations were observed in the top leaf sections on day 2 and in the middle leaf sections on day 6, these changes were not significant.

Malate concentrations were significantly affected by leaf position (ANOVA, p < 0.05) and the interaction between sampling time and leaf position (p < 0.05; Figure 4 H). Malate concentrations varied substantially across sampling times, with a particularly pronounced increase in the apical leaf sections toward the end of the experiment. Furthermore, malate concentrations at the apical leaf sections on day 12 were significantly higher than those in the middle leaf sections on days 6 (p = 0.0063) and 8 (p = 0.0101). These findings support a leaf-position-dependent pattern of malate accumulation, characterized by elevated concentrations in the apical leaf sections during the rewatering phase.

In summary, the combined analysis of chlorophyll, soluble sugars, and organic acids revealed distinct temporal and spatial patterns in leaf metabolism (Figure 4). Chlorophyll concentrations were primarily influenced by leaf position, with significantly higher concentrations in the apical leaf sections compared with the middle and basal leaf sections. In contrast, soluble sugar concentrations exhibited pronounced temporal changes. The temporal separation between sucrose and hexose accumulation indicated that these carbohydrates exhibited different accumulation dynamics during the experiment. Organic acid concentrations showed more heterogeneous responses. Citrate concentrations decreased significantly relative to the initial sampling point, whereas fumarate concentrations did not exhibit statistically significant changes. Malate concentrations showed a significant interaction between sampling time and leaf position, with a particularly pronounced increase during the rewatering phase at the apical leaf sections.

A bilayer neural network with sigmoid activation functions was used to assess associations between the measured variables and treatment phase, sampling day and leaf position. The bilayer neural network classifier showed good discriminatory performance across the three classification tasks which were (i) treatment classification (Figure 5 A), (ii) classification of sampling day (Figure 5 B), and (iii) classification of the leaf position (Figure 5 C). For treatment classification (Figure 5 A), the model achieved AUC values of 0.8433 for heat, 0.843 for heat/drought, and 0.7061 for rewatering phases, indicating lower classification performance for the rewatering condition. Classification of the experimental time course resulted in AUC values ranging from 0.661 to 0.887 (Figure 5 B), with the highest performance observed for 0d, i.e., initial conditions, (AUC = 0.887) and the lowest for 8d, i.e., the drought phase (AUC = 0.661). For the positional classes (Figure 5 C), classification was most accurate for the top position (AUC = 0.854), followed by the bottom (AUC = 0.750) and middle positions (AUC = 0.628). ANOVA-based feature association analysis revealed distinct sets of variables contributing to the separation of the respective classes. Treatment classification was primarily associated with citrate and sucrose, followed by glucose and malate, whereas classification of the (daily) time course was dominated by sucrose and glucose. In contrast, leaf positional classification was predominantly associated with chlorophyll content, followed by glucose (Figure 5 C, F). Overall, these results indicate that citrate and sucrose are good indicators for heat and drought effects while chlorophyll and glucose concentrations robustly predict leaf positions.

**Figure 5.**
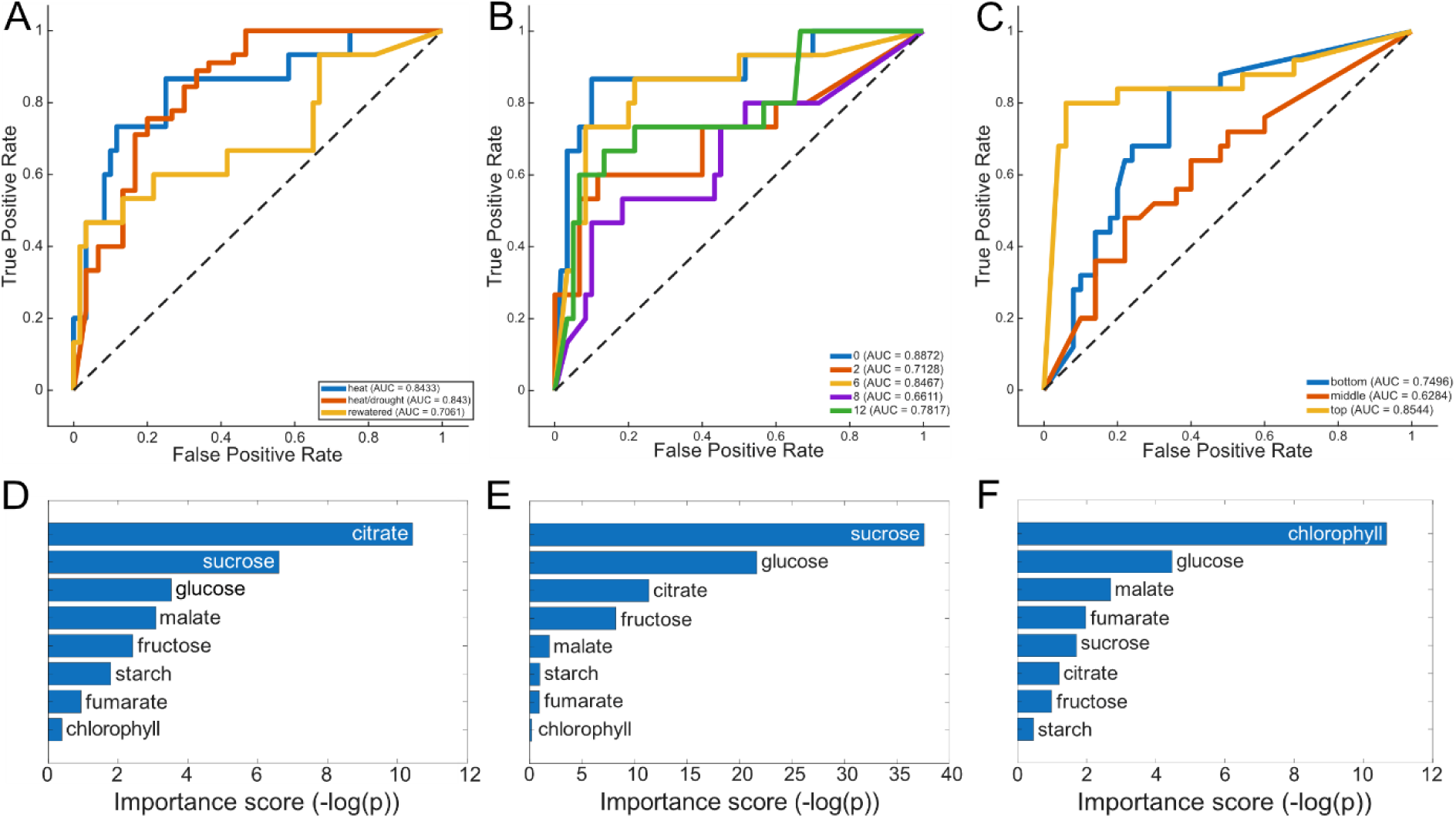
Classification performance and feature association of the bilayer neural network classifier. Receiver operating characteristic (ROC) curves and corresponding feature association scores are shown for the classification of (**A**) treatment (blue: heat, orange: heat/drought, yellow: rewatering), (**B**) time course of the drought/rewatering experiment in days (blue: 0d, orange: 2d, yellow: 6d, purple: 8d, green: 12d), and (**C**) leaf position (blue: bottom, orange: middle, yellow: top). ROC curves illustrate the discriminatory performance of the classifier for each class, with the corresponding area under the curve (AUC) reported in the plots. Feature importance scores (**D – F**) were determined using ANOVA and indicate the strength of univariate associations between individual physiological and metabolic variables and class labels. Higher importance scores, which represent the -log(p) – values of the ANOVA, indicate a stronger univariate association with the class labels. (**D**) feature association scores for classification of growth conditions, (**E**) feature association scores for classification of time course, (**F**) feature association scores for classification of leaf position.

## Discussion

### Drought response is spatially heterogeneous and delayed in apical leaf areas of *Sansevieria cylindrica*

The present study demonstrates that heat and drought exposure induce spatially heterogeneous physiological responses along the leaf blade of *Sansevieria cylindrica*. Most notably, the FW/DW ratio decreased significantly in basal leaf regions after eight days of water deprivation. Middle regions exhibited a similar but statistically non-significant tendency while apical regions remained largely unaffected. Following four days of rewatering, FW/DW ratios in basal and middle regions shifted towards initial heat values, while apical tissues exhibited a significant reduction. These findings indicate that the temporal dynamics of drought-associated changes in leaf tissue composition differ considerably along the longitudinal leaf axis. One possible explanation for this heterogeneous response lies in the specialized anatomical organization of *Sansevieria* leaves. The leaf mesophyll has been described to be differentiated into photosynthetically active chlorenchyma and internal water-storage tissues (Koller and Rost, 1988). The internal parenchyma of *Sansevieria* consists of regularly arranged living and dead cells, with the latter exhibiting specialized cell-wall structures and functioning as water reservoirs (Alfani et al., 1989). The authors observed that detached leaves lose water slowly and can rapidly recover which provides evidence for substantial hydraulic buffering capacity in this genus. Such anatomical properties could contribute to the maintenance of comparatively stable FW/DW ratios in apical tissues during drought, even as water availability declined. However, the findings of Alfani et al. (1989) also challenge a straightforward anatomical explanation for the delayed apical response. Although dehydration patterns were associated with leaf morphology, the authors found no relationship between the presence of specialized wall thickenings in dead parenchyma cells and the patterns of dehydration or rehydration. Consequently, the delayed FW/DW decrease observed in the present study cannot be attributed directly to these cell-wall structures. Instead, spatial differences in the relative abundance of storage tissues, hydraulic connectivity, or water allocation between living and non-living cells may be more relevant. Distinguishing between these mechanisms would require anatomical and hydraulic measurements at individual leaf positions.

The delayed apical response is also consistent with the broader concept that physiological responses to drought may lag behind changes in environmental water availability. Brodrick et al. (2019) demonstrated considerable spatial heterogeneity in forest drought resistance and showed that reductions in canopy water content can persist after meteorological drought conditions have subsided. Although their study addressed drought responses at a substantially larger spatial and temporal scale, it illustrates that water availability and physiological recovery are not necessarily synchronized. Similarly, drought responses involve coordinated hydraulic, metabolic, and regulatory processes rather than a single, immediately reversible response (Gupta et al., 2020). More recently, complex physiological, biochemical, and molecular adjustments were discussed in context of and associated with drought and rehydration concluding that rewatering does not necessarily restore all physiological parameters simultaneously (Song et al., 2026).

### Interactions between succulence, CAM metabolism, and spatial water conservation

The relationship between tissue hydration and metabolic activity is particularly complex in CAM plants because carbon assimilation and water conservation are temporally separated. Borland and Taybi (2004) described how CAM depends on coordinated diel changes in carbohydrate metabolism, organic acid accumulation, and circadian regulation. During the night, carbohydrate-derived phosphoenolpyruvate supports CO₂ fixation and the accumulation of organic acids, predominantly malate, whereas daytime decarboxylation provides internally concentrated CO₂ for photosynthesis. This metabolic organization allows CAM plants to restrict daytime stomatal opening and reduce transpirational water loss. Based on this mechanism, one might hypothesize that regions with higher CAM activity or carbohydrate availability exhibit enhanced drought resistance and therefore maintain their FW/DW ratios for longer periods. The relatively stable apical FW/DW during drought could be compatible with such a relationship, particularly because upper leaf regions received substantially higher photosynthetically active radiation than middle and basal regions. Greater light availability may support increased carbon assimilation and carbohydrate production, potentially influencing the capacity for osmotic adjustment and CAM-related carbon turnover.

However, Martin and colleagues provided important evidence against assuming a direct positive relationship between the amount of water-storage tissue and CAM activity (Martin et al. 2019). In their comparative investigation of ten *Sansevieria* species, CAM activity was negatively associated with leaf thickness and the hydrenchyma-to-chlorenchyma ratio, while no relationship was detected with mesophyll succulence. Their results suggest that CAM activity is more closely associated with the characteristics of photosynthetically active cells than with the overall volume of specialized water-storage tissue. Thus, a large water reservoir does not necessarily indicate greater CAM activity or a stronger metabolic contribution to drought tolerance. This distinction is particularly relevant for interpreting the present results. The delayed FW/DW response in apical tissues may reflect hydraulic buffering by specialized storage compartments without requiring enhanced local CAM activity. Conversely, metabolic differences between leaf regions may arise from differences in chlorenchyma abundance, light exposure, or carbon allocation independently of tissue hydration. The observed positional importance of chlorophyll content in the classification analysis supports the possibility that photosynthetic tissue characteristics contribute substantially to the spatial differentiation of the leaf.

### Carbohydrate and organic acid metabolism as potential indicators of drought-induced physiological changes

The temporal patterns in Figure 4 should be interpreted in the context of the treatment sequence shown in Supplementary Figure SF4: water was withheld until day 8, whereas elevated temperatures (30–33 °C during the day and 20–22 °C at night) were maintained throughout the experiment, including the four-day rewatering period. Thus, the changes observed before rewatering could reflect responses to progressive water limitation under sustained heat, where day 12 could represent a recovery of water supply under continued heat exposure. The transient sucrose accumulation on day 6, particularly in middle and basal tissues, followed by declining sucrose and maximal glucose and fructose concentrations on day 8, is consistent with a shift in carbohydrate partitioning as the combined treatment progressed. Sucrose cleavage and changes in carbon utilization could contribute to this sequence. Invertases may contribute by hydrolysing sucrose into glucose and fructose, thereby modifying carbon availability and sugar signalling; experimental studies in tomato have linked enhanced cell-wall invertase activity to improved drought responses and maintenance of fruit set under heat stress (Albacete et al., 2015; Liu et al., 2016). Thus, future studies might systematically reveal if (cell-wall) invertases also contribute to drought-induced metabolic adjustments in drought-adapted CAM species, such as *Sansevieria cylindrica*.

The increase in apical malate on day 12 relative to day 6 suggests a position-dependent adjustment of organic acid metabolism during rewatering under continued heat; because samples were collected at midday, this by itself is unlikely to reflect recovery or enhancement of nocturnal CAM activity. The early decrease in citrate, together with the absence of significant temporal effects on chlorophyll, starch and fumarate, further indicates a selective metabolic response rather than a uniform change across all measured pools. However, without separate heat-only and drought-only treatments, the individual effects of heat, drought and their interaction remain speculation.

Citrate and sucrose were the metabolites most strongly associated with treatment discrimination, followed by glucose and malate. Their importance suggests that drought-induced physiological changes are accompanied by adjustments in both carbohydrate allocation and organic acid metabolism. Geigenberger and colleagues demonstrated that carbon storage and partitioning are regulated through interactions between sugar availability, light-dependent redox signals, and metabolic enzyme activity (Geigenberger et al., 2005). Hence, these mechanisms provide a physiological basis for interpreting changes in soluble carbohydrates as indicators of altered metabolic regulation rather than merely passive consequences of water loss. Sucrose may be particularly relevant because it functions both as a transport carbohydrate and as a potential contributor to osmotic adjustment. Changes in sucrose availability could influence cellular osmotic potential, thereby contributing to water retention in metabolically active tissues. Glucose may similarly reflect altered carbohydrate turnover, mobilization of stored carbon, or changes in source–sink relationships. However, an increase in soluble sugar concentration would not necessarily demonstrate active osmotic adjustment. Higher sugar concentrations may reflect reduced carbon demand for growth or altered carbon utilization, while tissue dehydration can additionally increase concentrations through passive concentration of existing solute pools (Hummel et al., 2010; Turner, 2018). Therefore, the importance of sucrose and glucose for classification supports their association with drought-related metabolic changes but does not establish a causal contribution to water retention.

The prominence of citrate in treatment classification is particularly interesting because malate is generally considered the principal organic acid associated with nocturnal carbon fixation in CAM plants. Borland and Taybi (2004) emphasized the reciprocal diel fluctuations of carbohydrates and organic acids and their regulatory importance for CAM. Nevertheless, the comparatively high discriminatory importance of citrate suggests that the metabolic response to drought may extend beyond the canonical malate-centered CAM cycle. Citrate may reflect changes in respiratory carbon metabolism, organic acid storage, or broader adjustments in cellular carbon balance.

An alternative explanation is that citrate and sucrose are particularly effective treatment discriminators because their concentrations respond to multiple physiological processes simultaneously. Their statistical importance does not necessarily indicate that they are the principal regulators of drought tolerance. Furthermore, because metabolite concentrations were determined at midday, the present measurements capture only one phase of the diel CAM cycle. Differences in metabolite abundance could therefore reflect changes in the timing or amplitude of metabolic oscillations rather than changes in total daily metabolic flux. Resolving this distinction would, for example, require measurements throughout the day–night cycle.

### Metabolic signatures of stress persistence and incomplete recovery

The relatively low classification accuracy of the rewatering phase may provide additional insight into the delayed FW/DW response of apical tissues. Following rewatering, basal and middle regions exhibited a tendency towards restoration of their initial FW/DW ratios, whereas apical regions showed a significant decrease. This indicates that samples collected during recovery did not necessarily represent a uniform physiological state. Instead, different leaf regions may have undergone distinct stages of recovery or continued to exhibit physiological consequences of the preceding drought period. Such an interpretation is compatible with the concept of stress-induced metabolic persistence. It has been demonstrated before that *Sansevieria trifasciata* can retain physiological effects of previous exposure to particulate matter and volatile organic compounds after a recovery period (Permana et al., 2024). Their metabolomic analyses identified the accumulation of ornithine-related compounds, and exogenous ornithine application improved stress tolerance through effects associated with proline accumulation and antioxidant activity. These findings provide evidence that metabolic adjustments in *Sansevieria* can persist beyond the original stress exposure and influence subsequent physiological responses.

Although the stressors investigated by Permana and colleagues differ fundamentally from drought (Permana et al., 2024), their results support the broader possibility that metabolic states in *Sansevieria* may retain information about previous environmental conditions. Accordingly, the delayed apical FW/DW decrease observed after rewatering could reflect physiological processes initiated during drought that remain active during rewatering, i.e., recovery phases. However, the present data does not yet establish a stress-memory mechanism. Demonstrating such mechanism would require evidence that prior drought exposure alters the response to a subsequent stress event, preferably together with measurements of persistent metabolic or regulatory changes, as e.g. shown for gene regulatory networks (Deng et al., 2026).

An alternative interpretation is that the recovery phase represents a transient mixture of physiological states rather than a distinct metabolic condition. Some tissues may already have restored water balance, whereas others may continue to undergo changes in carbohydrate turnover, organic acid metabolism, or photosynthetic performance. This could explain why recovery samples were less accurately classified than (initial) heat and drought samples. Importantly, however, lower classification accuracy alone does not prove physiological heterogeneity, because it may also arise from limited sample size, overlapping metabolite distributions, or model uncertainty.

### Photosynthetic responses may precede detectable changes in apical FW/DW

The chlorophyll fluorescence measurements provide additional evidence that physiological stress responses are not necessarily synchronized with changes in FW/DW. During drought exposure, the effective quantum yield of photosystem II, Y(II), decreased significantly between days 6 and 8, while non-regulated energy dissipation, Y(NO), increased. In contrast, Y(NPQ) exhibited no significant systematic changes. Because chlorophyll fluorescence was measured near the leaf tip, these observations are particularly relevant to the apical FW/DW response.

The decrease in Y(II) despite the absence of a significant apical FW/DW reduction during drought suggests that photosynthetic performance may respond to stress before corresponding changes in tissue mass ratios become detectable. This interpretation is consistent with the broader physiological framework described by Gupta et al. (2020), in which photosynthetic inhibition and metabolic adjustments may arise through stomatal and non-stomatal limitations during water deprivation.

However, chlorophyll fluorescence parameters do not necessarily respond uniformly to drought. Zait and colleagues investigated the dynamic responses of fluorescence parameters across six plant species and demonstrated that different indicators of photosynthetic performance respond at different stages of drought progression (Zait et al., 2024). In particular, maximal NPQ and Y(NPQ) declined when Fv/Fm fell below approximately 0.7, while other fluorescence parameters showed different response patterns. Their findings indicate that the interpretation of individual fluorescence parameters depends on stress severity and measurement conditions. This is important for the present study because the absence of significant changes in Y(NPQ) should not be interpreted as evidence that regulated photoprotection was unaffected by drought. Rather, the observed fluorescence response may reflect the particular stress intensity, light environment, or temporal stage of the experiment.

The simultaneous decline in Y(II) and increase in Y(NO) indicates that photochemical energy utilization decreased while non-regulated energy dissipation became more prominent. These changes may reflect an increasing imbalance between absorbed light energy and its utilization in photosynthetic metabolism. However, because Y(II), Y(NPQ), and Y(NO) represent complementary fractions of absorbed excitation energy, their negative associations are not statistically independent. Consequently, the strong inverse relationship between Y(II) and Y(NO) should be interpreted primarily as a redistribution of excitation-energy partitioning rather than independent evidence for a specific mechanism of photoinhibition.

Taken together, the chlorophyll fluorescence and FW/DW results support the hypothesis that functional photosynthetic changes can precede significant alterations in apical tissue mass ratios. Nevertheless, it remains speculation here whether the fluorescence response is driven primarily by declining water availability, altered carbon metabolism, temperature stress, or interactions among these factors.

### Metabolic profiles as indicators of physiological state and potential predictors of stress responses in CAM plants

The neural network classification results highlight the potential of metabolic and physiological profiles to infer stress-associated states in *Sansevieria cylindrica*. Their ability to discriminate experimental phases, sampling times and leaf positions suggests that these profiles capture coordinated responses to environmental conditions and spatial organization within the leaf. This provides a promising basis for developing metabolic biomarkers that identify plant physiological status and, with temporal validation, may anticipate subsequent responses to stress and rewatering. The distinctive apical profile is particularly informative because these tissues exhibited a delayed reduction in FW/DW. This correspondence raises the possibility that biochemical signatures could help identify tissues undergoing physiological adjustment before changes in tissue hydration become detectable. Chlorophyll contributed substantially to positional discrimination, suggesting that combining metabolic markers with indicators of photosynthetic tissue composition may improve state estimation. The partial independence of photosynthetic tissue characteristics and water-storage capacity reported earlier further supports integrating complementary measurements to capture different dimensions of succulent leaf function (Martin et al., 2019).

## Conclusion

Based on our findings, we suggest that spatial gradients in leaf development, physiology, and metabolism provide a natural framework for investigating plant–environment interactions and the mechanisms underlying stress acclimation and recovery (Figure 6). Combining spatially resolved biochemical and physiological measurements with experimental time courses and environmental treatments enables the disentanglement of developmental processes from stress-induced responses and the identification of spatiotemporal patterns of metabolic adjustment. Integrating these multidimensional datasets through statistical analysis and predictive modelling could further extend the classification of physiological states towards forecasting stress responses and recovery trajectories. A promising next step would be to test whether biochemical profiles measured during heat exposure or early drought predict subsequent changes in FW/DW, photosynthetic performance, or recovery following rewatering. In this context, *S. cylindrica* represents a valuable model for investigating how spatially resolved biochemical signatures reflect and potentially predict plant responses to environmental stress. Given the central roles of carbohydrate turnover and organic acid storage in CAM metabolism, similar approaches may also be applicable to other CAM species, accounting for differences in anatomy and metabolic regulation. Finally, validation across stress regimes and species will be essential to establish the robustness, predictive capacity, and transferability of stress markers, ultimately advancing mechanistic understanding of plant–environment interactions, stress acclimation and adaptation.

**Figure 6.**
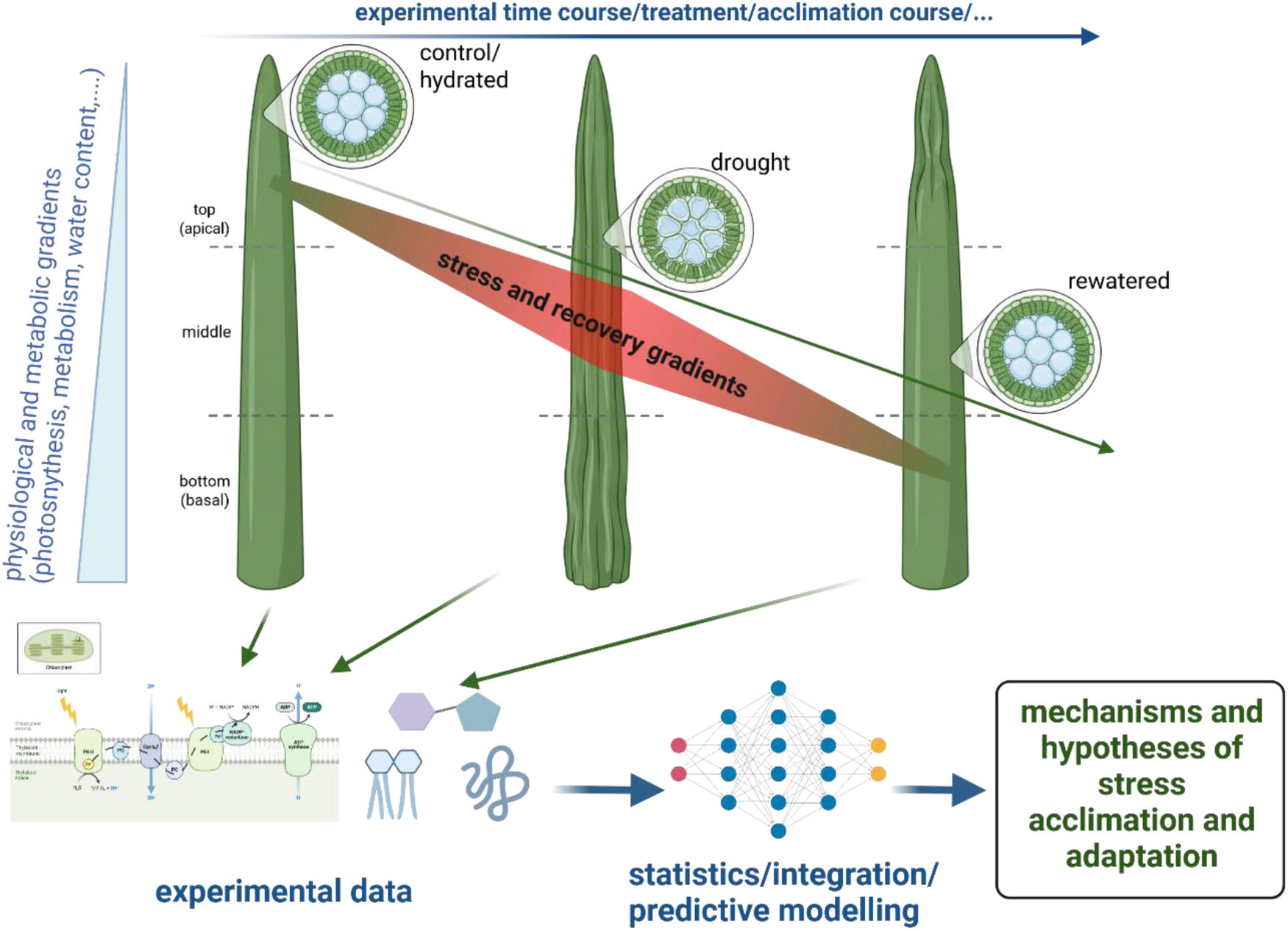
Exploiting spatial gradients in leaf physiology and metabolism to decode plant–environment interactions. Spatial gradients in leaf development, physiology, and metabolism, combined with experimental time courses and environmental treatments, provide a framework for distinguishing developmental variation from stress-induced responses and tracking stress acclimation and recovery. Integration of spatially resolved physiological, biochemical, and molecular data through statistical analysis and predictive modelling facilitates the identification of stress-associated signatures, regulatory relationships, and potential mechanisms underlying plant acclimation and adaptation to environmental change. Created in BioRender. Naegele, T. (2026) https://BioRender.com/ub828ku

## Acknowledgements

We thank the members of Plant Evolutionary Cell Biology at LMU München, and particularly Vladimir Brodsky, for advice and help with experimental protocols. The authors acknowledge the use of ChatGPT powered by GPT-6 to support creation of the abstract and to suggest reorganization of Introduction and Discussion section for clarity and flow. All AI-generated suggestions were reviewed, revised, and approved by the authors, who take full responsibility for the accuracy and integrity of the work.

## Data availability statement

Data of this study is available in the supplements.

## Author contributions

<u>RPV:</u> Performed experiments, statistics, and data evaluation. <u>TN:</u> conceived the study, performed statistics and data evaluation, and wrote the manuscript.

## Supplementary Tables

**Supplementary Table ST1. Chlorophyll fluorescence measurements during heat exposure, combined heat and drought, and rewatering under continued heat in *Sansevieria cylindrica*.** Individual pulse amplitude modulation (PAM) measurements are provided for experimental days 0, 2, 6, 8 and 12, together with clock time (TimeOfDay) and the corresponding light or dark period (DayNight; day: 06:00–18:00; night: 18:00–06:00). Y(II) denotes the effective quantum yield of photosystem II, Y(NPQ) the quantum yield of regulated non-photochemical energy dissipation, and Y(NO) the quantum yield of non-regulated energy dissipation. All yields are dimensionless. Measurements were recorded near the leaf tip.

**Supplementary Table ST2. Spatially resolved carbohydrate, organic acid and chlorophyll measurements in *Sansevieria cylindrica*.** Data is provided for apical (Top), middle (Middle) and basal (Bottom) leaf sections sampled at midday on experimental days 0, 2, 6, 8 and 12. Sucrose, glucose, fructose, malate, fumarate and citrate concentrations are expressed as µmol g⁻¹ fresh weight; starch is expressed as µmol C6 equivalents g⁻¹ fresh weight. Chlorophyll is reported as µg gFW⁻¹.

**Supplementary Table ST3. Two-way ANOVA results for chlorophyll and metabolite concentrations in *Sansevieria cylindrica*.** The effects of sampling day (Day), leaf position (Leaf_position) and their interaction (Day:Leaf_position) were evaluated for chlorophyll, sucrose, starch, glucose, fructose, citrate, fumarate and malate. p_Raw denotes unadjusted p-values, and q_BH denotes p-values adjusted using the Benjamini–Hochberg procedure to control the false discovery rate. Significant_Raw and Significant_BH indicate significance before and after adjustment, respectively: ns, p or q ≥ 0.05; *, < 0.05; **, < 0.01; ***, < 0.001.

**Supplementary Table ST4. Tukey–Kramer post hoc comparisons of chlorophyll and metabolite concentrations in *Sansevieria cylindrica*.** Pairwise comparisons are provided for sampling days (Day), leaf positions (Leaf_position) and combinations of sampling day and leaf position (Day × Leaf_position). Variable identifies the measured compound; chloro_gFW denotes chlorophyll concentration expressed per fresh weight. Group1 and Group2 identify the compared groups, Difference reports the estimated difference between group means, and LowerCI and UpperCI indicate the corresponding confidence limits. p_Adjusted denotes the p-value adjusted for multiple comparisons using the Tukey–Kramer procedure within each comparison family. Significance is indicated as ns, p ≥ 0.05; *, p < 0.05; **, p < 0.01; ***, p < 0.001.

## Supplementary Figures

**Supplementary Figure SF1.**
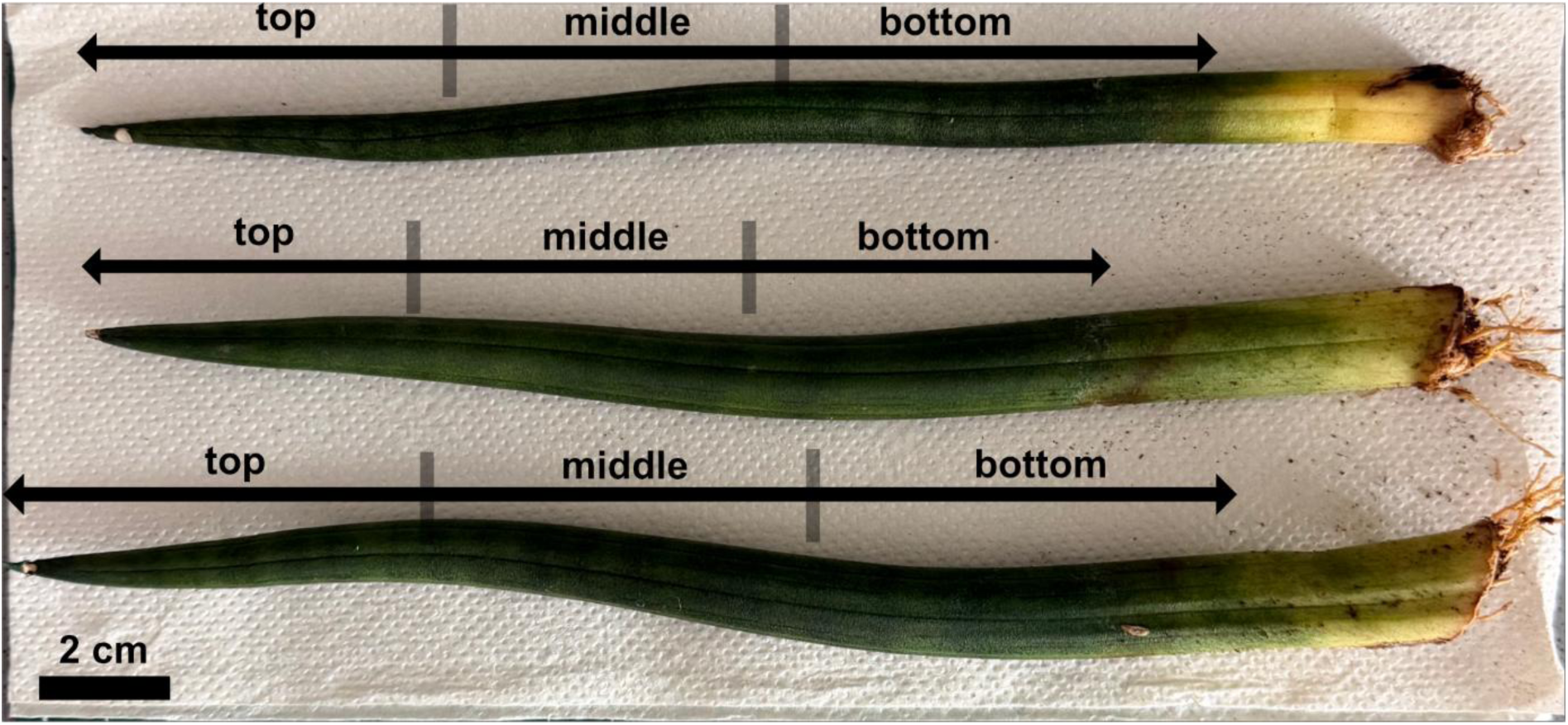
Leaves and sections for analysis of *Sansevieria cylindrica*. Arrows indicate the above-ground part of each leaf. The sections top, middle and bottom were defined by 1/3 of the total aboveground leaf length for each leaf individually. The size bar (bottom left) represents 2 cm.

**Supplementary Figure SF2.**
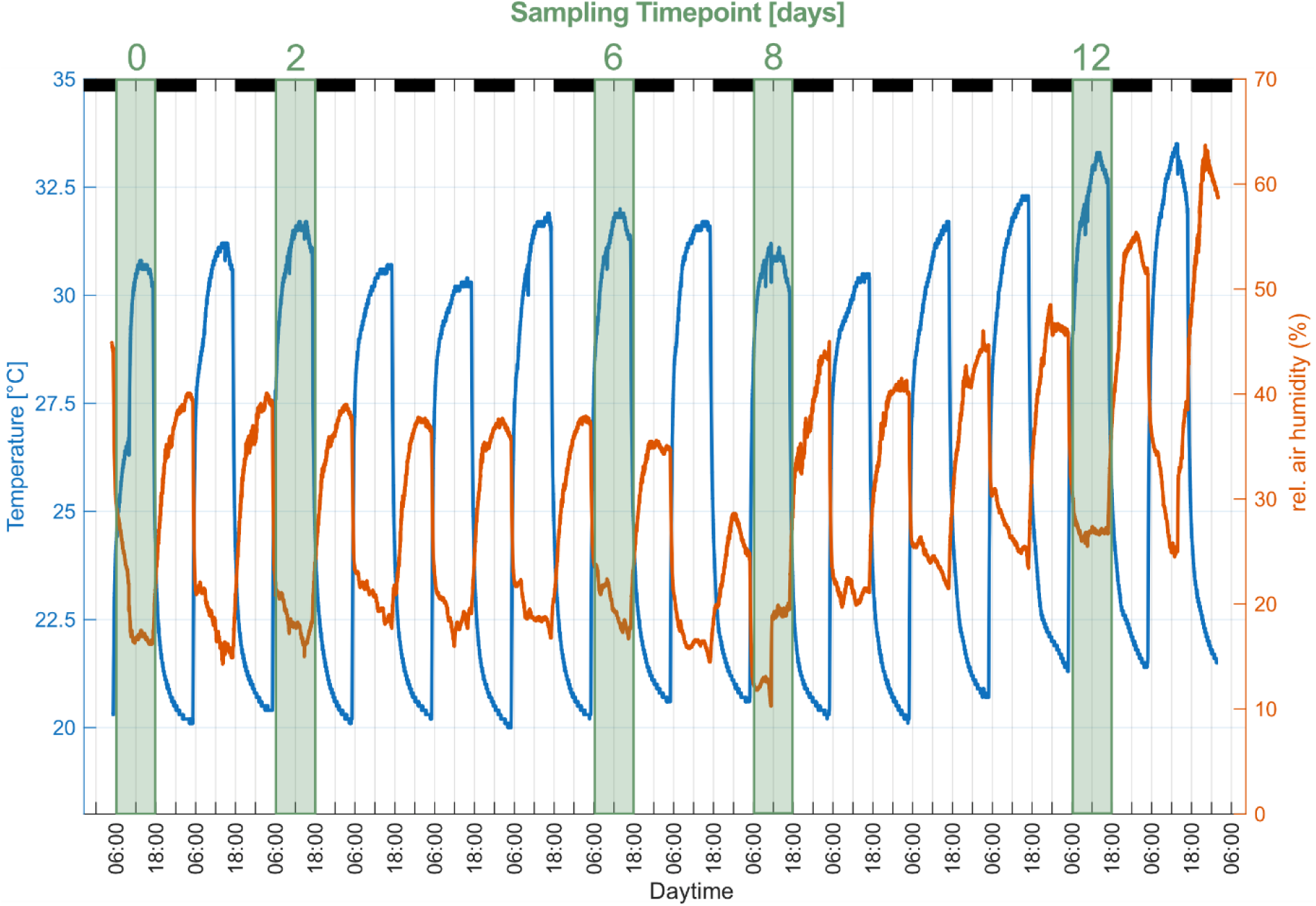
Air temperature and air humidity during the experimental time course. The time course of air temperature is shown by the blue line [°C], relative air humidity (in %) by the orange line. Sampling time points (in days) are indicated by green boxes and labelling on the top. Time labels on the bottom show the time of day.

**Supplementary Figure SF3.**
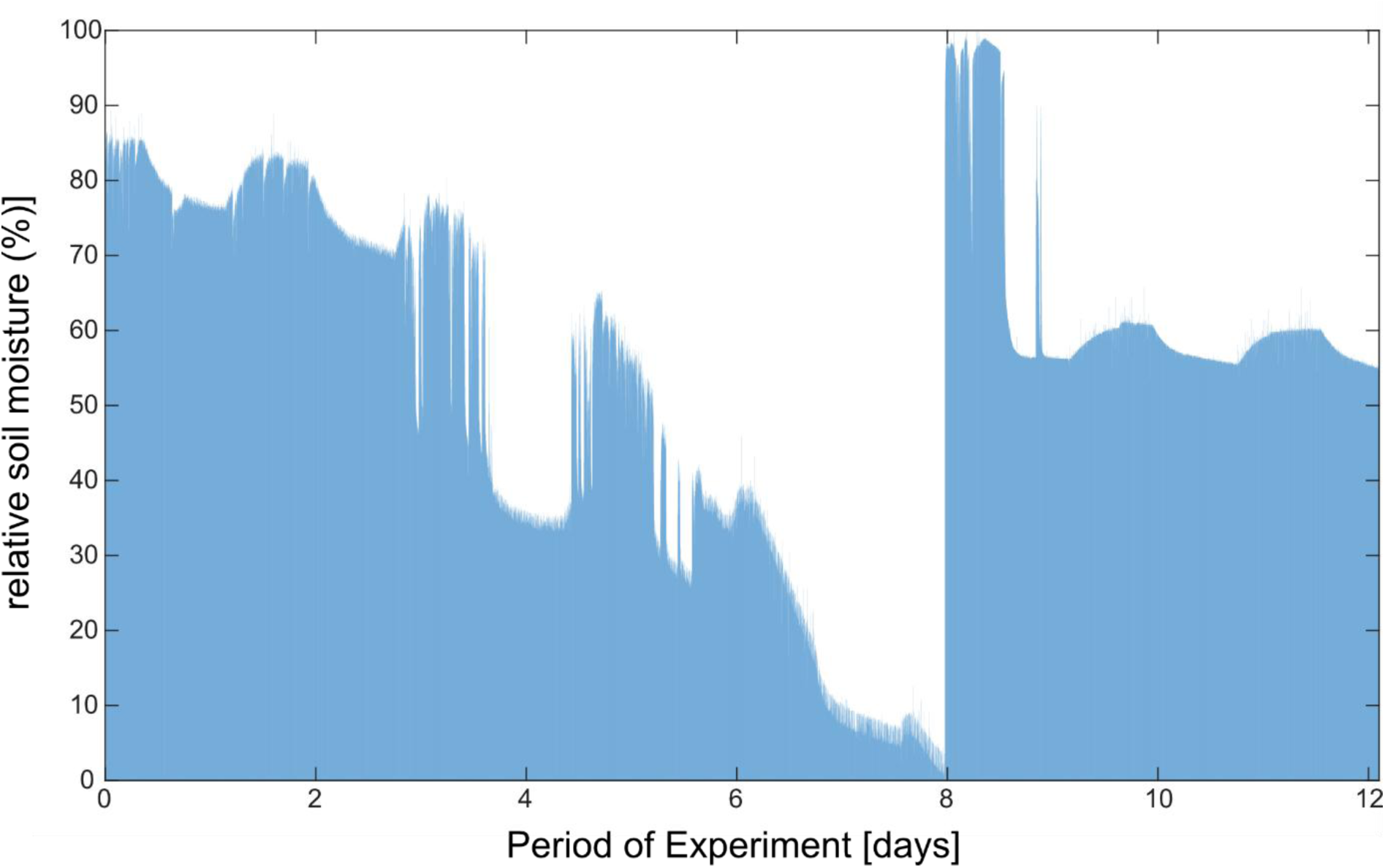
Time course of relative soil moisture during the experiment. Wet soil was defined by relative soil moisture between 100 and 75 %, humid soil between 74 and 50 %, and dry soil by values < 50 %.

**Supplementary Figure SF4.**
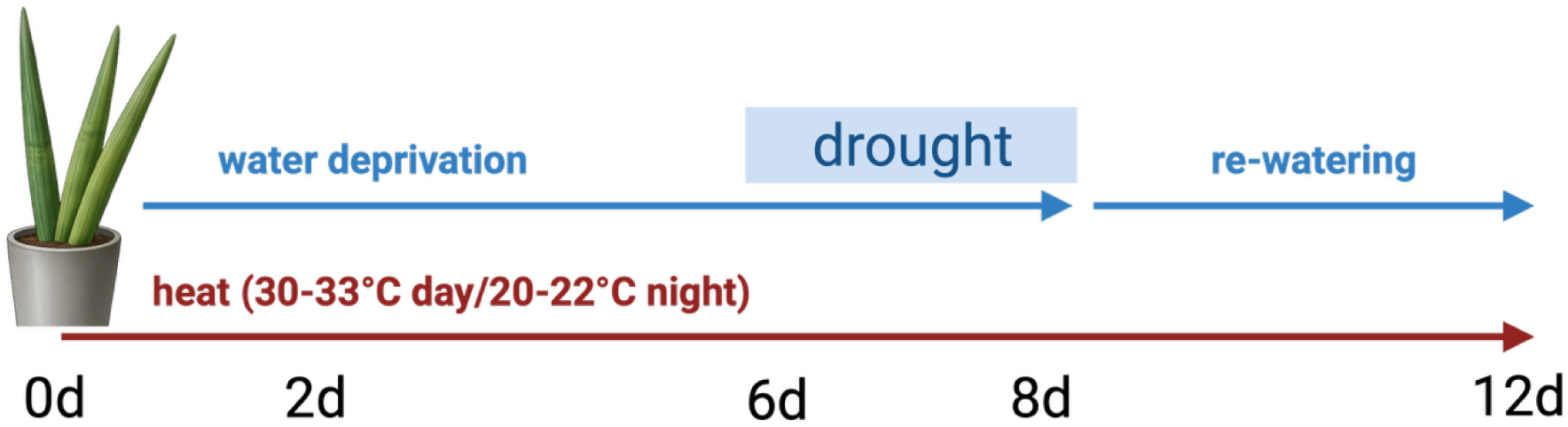
Experimental setup and sampling time course. *Sansevieria cylindrica* plants were exposed to heat throughout the 12-day experiment (30–33 °C during the day and 20–22 °C at night). Water was withheld from day 0 to day 8, comprising an initial heat phase (days 0–6) followed by combined heat and drought (days 6–8). After sampling on day 8, plants were rewatered and maintained under continued heat until day 12. Samples were collected on days 0, 2, 6, 8 and 12. Blue arrows indicate water deprivation and rewatering; the red arrow indicates continuous heat exposure. Created in BioRender. Naegele, T. (2026) https://BioRender.com/yklhvse.

